# *Smad4* is essential for epiblast scaling and morphogenesis after implantation, but nonessential prior to implantation in the mouse

**DOI:** 10.1101/2024.01.23.576717

**Authors:** Robin E. Kruger, Tristan Frum, A. Sophie Brumm, Stephanie L. Hickey, Kathy K. Niakan, Farina Aziz, Marcelio A. Shammami, Jada G. Roberts, Amy Ralston

## Abstract

Bone Morphogenic Protein (BMP) signaling plays an essential and highly conserved role in axial patterning in embryos of many externally developing animal species. However, in mammalian embryos, which develop inside the mother, early development includes an additional stage known as preimplantation. During preimplantation, the epiblast lineage is segregated from the extraembryonic lineages that enable implantation and development *in utero*. Yet, the requirement for BMP signaling in mouse preimplantation is imprecisely defined. We show that, in contrast to prior reports, BMP signaling (as reported by SMAD1/5/9 phosphorylation) is not detectable until implantation, when it is detected in the primitive endoderm – an extraembryonic lineage. Moreover, preimplantation development appears normal following deletion of maternal and zygotic *Smad4,* an essential effector of BMP signaling. In fact, mice lacking maternal *Smad4* are viable. Finally, we uncover a new requirement for zygotic *Smad4* in epiblast scaling and cavitation immediately after implantation, via a mechanism involving FGFR/ERK attenuation. Altogether, our results demonstrate no role for BMP4/SMAD4 in the first lineage decisions during mouse development. Rather, multi-pathway signaling among embryonic and extraembryonic cell types drives epiblast morphogenesis post-implantation.

**Summary Statement:** Gene expression, gene deletion, and pathway visualization evidence show that *Smad4*-dependent signaling is first active after mouse embryo implantation, when it promotes epiblast morphogenesis non-cell autonomously.

## Introduction

In animal embryos, including mice, frogs, fish, and flies, the Bone Morphogenic Protein (BMP) signaling pathway oversees critical patterning events early in development. In non-mammalian species, BMP signaling is critical for specification of the dorsal/ventral axis of the early embryo (De Robertis and Sasai, 1996; O’Connor et al., 2006; Zinski et al., 2018). However, the mammalian embryo has an additional developmental task immediately following fertilization: specification of the extraembryonic lineages that will give rise to placenta and yolk sac and enable development within the mother. Published studies support roles for BMP signaling in both extraembryonic lineage specification, prior to implantation, and subsequent axial patterning, which occurs after implantation. However, differences in technical approaches used, as well as challenges intrinsic to mouse, have limited origination of a universally accepted model of the role of BMP signaling in mouse embryos throughout pre- and post-implantation stages.

BMP is one of several related and highly conserved molecular signaling pathways belonging to the Transforming Growth Factor beta (TGFβ) superfamily of cytokines. The molecular mechanisms of TGFβ signaling have been carefully studied (Chang, 2016; Massagué and Sheppard, 2023). BMP proteins, like other members of the TGFβ pathway, are secreted ligands that elicit cellular responses by binding to heterodimeric, transmembrane serine-threonine kinase receptors. The activated receptor complex then phosphorylates members of a family of intracellular effectors known as receptor-associated SMADs (r-SMADs). Phosphorylation of r-SMADs allows their association with a co-factor SMAD and accumulation in the nucleus, where they impact chromatin and transcription (Hill, 2016). In mammals, r-SMAD activity is encoded by several *Smad* paralogues, with SMAD1, SMAD5, and SMAD9 (also known as SMAD8) primarily transducing BMP signals and SMAD2 and SMAD3 primarily transducing Nodal, Activin, and TGFβ. Notably, the mammalian genome encodes a single co-Smad, SMAD4, which is shared by BMP, Nodal, Activin, and TGFβ signaling pathways.

Across species, BMP signaling has been visualized in embryos using antibodies that specifically recognize the phosphorylated form of the BMP-responsive r-SMAD(s). This approach has been used to observe gradients of BMP signaling activity that correspond with the dorsal/ventral axis in fly, fish, and frog embryos (Dorfman and Shilo, 2001; Plouhinec and De Robertis, 2009; Schohl and Fagotto, 2002; Tucker et al., 2008). In mouse, no graded pSMAD1/5/9 pattern has been reported. Prior to implantation, pSMAD1/5/9 is reportedly detected in all cell types of the embryo at multiple stages (Graham et al., 2014; Reyes de Mochel et al., 2015). After implantation, pSMAD1/5/9 is detected within a subdomain of extraembryonic cells, and not within the embryo itself until it is detected in primordial germ cells and emerging mesoderm during gastrulation (Senft et al., 2019). These observations suggest fundamental differences in the roles of BMP signaling between mammalian and non-mammalian animal embryos, but raise the need for additional, functional lines of evidence.

In mice, individual members of the BMP signaling pathway appear to be dispensable prior to embryonic day 6.5 (E6.5). Knockout of genes encoding the predominant ligand *Bmp4* (Lawson et al., 1999; Winnier et al., 1995), the receptors *Bmpr2* (Beppu et al., 2000)*, Bmpr1a* (Mishina et al., 1995)*, Actr1a* (Gu et al., 1999), the r-SMADs encoded by *Smad1* (Tremblay et al., 2001) and *Smad5* (Chang et al., 1999), and the co-Smad *Smad4* (Sirard et al., 1998; Yang et al., 1998; Yang et al., 2002) all point to essential roles for BMP signaling in extraembryonic mesoderm, extraembryonic endoderm, and germ cell development. Mechanistically, BMP also interacts with Nodal to pattern the visceral endoderm and to specify distal, and then anterior visceral endoderm, structures required to spatially pattern the embryo and specify the primitive streak (Robertson, 2014; Waldrip et al., 1998; Yamamoto et al., 2009). These events define gastrulation and anterior/posterior axial patterning in mouse, processes which therefore rely on BMP signaling. None of these studies reported that BMP signaling loss-of-function had any effect on development prior to E5.5. However, maternal gene products provided within the oocyte could complicate interpretation of knockout phenotypes resulting from zygotic gene deletion only. Indeed, evidence exists that BMP pathway members are maternally supplied and functional in embryos of other animal species (Das et al., 1998; Faure et al., 2000; Kramer et al., 2002; Miyanaga et al., 2002; Zhang et al., 2020) Finally, mouse embryos are particularly challenging to recover between E4.5 and E6.5 and we lack an *in vitro* protocol that robustly recapitulates *in vivo* development during these stages, presenting a barrier to the facile testing of a possible role for BMP signaling during the peri-implantation period.

By contrast, preimplantation embryos are relatively easy to isolate and culture *in vitro*. Accordingly, several studies have suggested a role for BMP signaling in preimplantation development. Culturing preimplantation embryos in the presence of small-molecule BMP inhibitors led to decreased numbers and cell cycle rate of extraembryonic trophectoderm (TE) and primitive endoderm (PrE) cells, as well as changes in expression of lineage-specific transcription factors, including markers of PrE (SOX17, GATA6), TE (CDX2), and inner cell mass (ICM, OCT4) (Graham et al., 2014; Reyes de Mochel et al., 2015; Stuart et al., 2019). Some of these observations were recapitulated following microinjection of siRNA against *Bmp4* or overexpression of dominant-negative forms of *Bmpr2* (Graham et al., 2014). Overexpression of dominant-negative *Smad4* reportedly phenocopied loss of the upstream signaling components. In principle, these approaches could interfere with the activities of both maternally and zygotically expressed signaling components and thereby achieve more extreme loss of function. However, pSMAD1/5/9 was not examined in these manipulated embryos, so the extent to which these manipulations disrupted BMP signaling is unclear. Moreover, inhibitors are prone to off-target effects, which could further confound interpretation of results (Lowery et al., 2016).

In the present study, we visualize pSMAD1/5/9 in wild-type and embryos in which *Bmp4* has been maternally and zygotically deleted, as well as lineage specification and morphogenesis in embryos lacking maternal and zygotic *Bmp4* or *Smad4* throughout preimplantation, peri-implantation, and early post-implantation stages. We report that, in contrast to previous studies, BMP signaling is apparently dispensable during mouse preimplantation development. However, we observed that SMAD4-mediated signaling is essential for peri-implantation epiblast morphogenesis shortly after implantation, when it helps attenuate FGF/ERK signaling to enable the timely execution of epiblast morphogenetic events.

## Results

### Phosphorylated SMAD1/5/9 is first detectable in peri-implantation embryos

To determine when BMP signaling becomes active in the mouse embryo, we first developed a method to examine the localization of transcription factors SMAD1, 5, and 9, which are phosphorylated in response to ligand/receptor binding (Dijke and Hill, 2004). To achieve this, we used immunofluorescence and an antibody that recognizes phosphorylated SMAD1/5/9 (pSMAD1/5/9, Fig. 1) (Senft et al., 2019; Xu et al., 2019; Yuan et al., 2015). We did not detect pSMAD1/5/9 in preimplantation embryos flushed from uteri between E3.75-E4.25 (Fig 1A, Supp. Fig. 1A). We first observed pSMAD1/5/9 in E4.5 peri-implantation embryos (Fig. 1A), when it was detected in nuclei of a few inner cell mass cells in 29% of embryos examined (Fig. 1B-C). By E4.75, when embryos have undergone implantation, we observed pSMAD1/5/9-positive cells in 87.5% of the embryos evaluated (Fig. 1A-C). Starting at E5.0, we observed pSMAD1/5/9-positive cells within 100% of embryos examined (Fig. 1A-C, Supp. Fig. 1A). The observed pSMAD1/5/9 overlapped with a sub-set of GATA6-expressing primitive endoderm (E4.5 and E4.75) and visceral endoderm (E5.5) cells (Fig. 1A).

**Figure 1.**
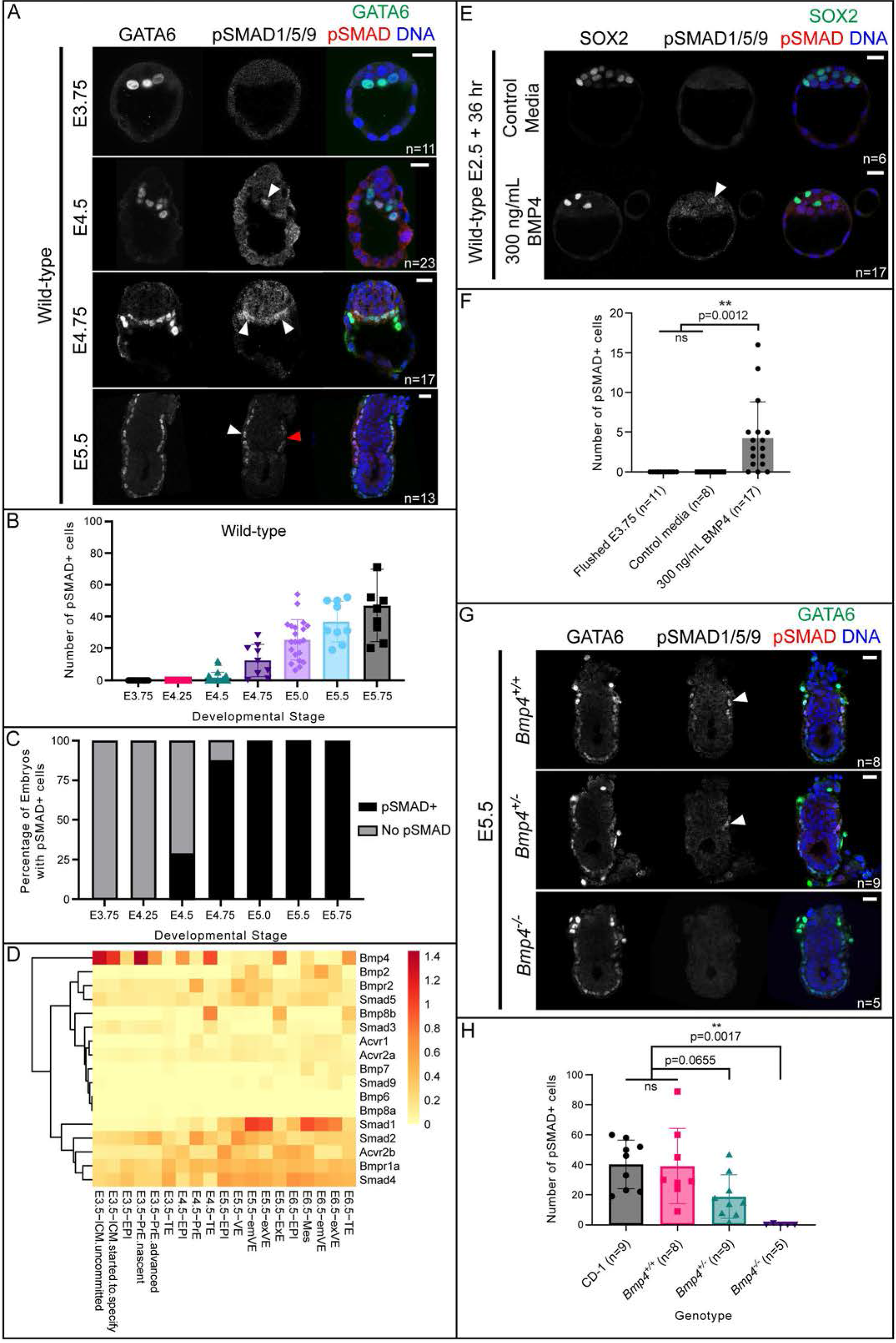
BMP signaling becomes active in primitive endoderm at implantation. A) SMAD1/5/9 phosphorylation (pSMAD1/5/9) in wild-type CD-1 embryos at E3.75, E4.5, E4.75, and E5.5. In all cases, positive pSMAD1/5/9 signal co-localizes with GATA6 as a marker of primitive endoderm and visceral endoderm. B) Quantification of total number of pSMAD1/5/9-positive cells in wild-type embryos in A and Supplemental Figure 1A. C) Quantification of the percentage of embryos from A and Supplemental Figure 1A which display any pSMAD1/5/9-positive cells versus no pSMAD1/5/9-positive cells. D) Heat map of the mean normalized expression of BMP pathway genes from scRNA-seq data from Nowotschin et al., 2019. E) pSMAD1/5/9 in wild-type embryos collected at E2.75 and cultured for 36 hours in media containing 300 ng/mL exogenous BMP4. F) Quantification of the total number of pSMAD1/5/9-positive cells in embryos from E revealed significantly more pSMAD1/5/9-positive cells in BMP4-treated embryos. G) pSMAD1/5/9 staining is absent in *Bmp4* z null embryos at E5.5. H) Quantification of total number of pSMAD1/5/9-positive cells in wild-type and *Bmp4*-null embryos at E5.5 revealed significantly fewer pSMAD1/5/9-positive cells in *Bmp4-*null embryos. All pairwise comparisons were assessed by analysis of variance (ANOVA) with Tukey’s post-hoc test. White arrowheads indicate positive pSMAD1/5/9 signal. Red arrowhead indicates a GATA6+ cell which does not express pSMAD1/5/9. Scale bars represent 10 μm.

To determine whether the observed pSMAD1/5/9 signal was specific, we treated E5.5 wild-type embryos with LDN-193189 (LDN hereafter) which has been used to disrupt BMP signaling in mouse embryos (Graham et al., 2014; Reyes de Mochel et al., 2015). A concentration of 1 µM LDN was reported as sufficient to inhibit BMP signaling in preimplantation mouse embryos (Reyes de Mochel et al., 2015). However, we found that treatment with 1 µM LDN was highly toxic to embryos (Supp. Fig. 1B). Nevertheless, treatment with 0.25 µM LDN led to complete loss of pSMAD1/5/9 signal in E5.5 embryos (Supp. Fig 1B). Altogether, these observations suggest that BMP signaling becomes active around the time of embryo implantation but is not active during preimplantation stages.

### BMP pathway members are present, but largely inactive, prior to implantation

A prior report showed that BMP4 is sufficient to influence gene expression in preimplantation mouse embryos CITE, suggesting that preimplantation embryos can respond to exogenous BMP signals. We therefore examined expression dynamics of genes encoding BMP pathway members during preimplantation stages. We analyzed published single-cell RNA-seq data from mouse embryos at stages E3.5-E6.5 (Nowotschin et al., 2019). At E3.5, many core components of canonical BMP signaling were detectable, including the ligand *Bmp4,* Type I receptor *Bmpr1a*, Type II receptors *Bmpr2* and *Acvr2b,* receptor-associated SMAD *Smad5,* and co-factor SMAD *Smad4* (Fig. 1D and Supp. Fig. 2).

Next, we investigated whether pSMAD1/5/9 could be induced in preimplantation embryos treated with exogenous BMP. We cultured compacted 8-cell stage embryos (E2.75) in 300 ng/ml BMP4 for 36 hours to the blastocyst stage (equivalent in cell number to E3.75, as confirmed by cell counts). Although we did not observe pSMAD1/5/9 in any control embryos cultured in unsupplemented medium, we observed low, but detectable levels of pSMAD1/5/9 in 82% (n=14/17) of embryos treated with exogenous BMP4, further supporting it faithful detection of pSMAD1/5/9 by immunofluorescence analysis (Fig. 1E-F). Notably, pSMAD1/5/9 was detected only in the ICM but did not preferentially colocalize with either SOX2-positive epiblast (EPI) or SOX2-negative PrE cells, indicative of normal ICM differentiation. Therefore, we conclude that BMP signaling is not highly active during preimplantation development, but ICM cells are competent to respond to exogenous BMP signals at these stages, consistent with published investigations CITE.

We next evaluated pSMAD1/5/9 in embryos shortly after implantation. Consistent with prior reports (Senft et al., 2019; Yamamoto et al., 2009), we detected pSMAD1/5/9 within a zone of the visceral endoderm (VE) that flanks the extraembryonic ectoderm (EXE) at E5.5 and E5.75 (Fig. 1A, Supp. Fig. 1A). This observation is also consistent with evidence that several key components, including *Bmp2, Smad1, Smad5,* and *Bmpr2,* are substantially upregulated around the time of implantation (E4.5-E5.5), particularly in the PrE/VE lineage (Fig. 1D, Supp. Fig 2A-B). Notably, culturing E5.5 embryos in the presence of exogenous BMP4 for 6 hours was sufficient to expand the zone of pSMAD1/5/9 within the VE in a dose-dependent manner (Supp. Fig. 1C). Thus, the availability of ligand could limit the extent of pathway activation, during both pre- and post-implantation stages.

Finally, we evaluated pSMAD1/5/9 in *Bmp4*-null embryos at E5.5. We were unable to detect pSMAD1/5/9in *Bmp4*-null embryos, although it was observed at wild-type levels and localization in homozygous wild-type littermate controls (Fig. 1G-H). In *Bmp4* heterozygous embryos, we observed an intermediate phenotype where some pSMAD1/5/9 was detectable but trended toward lower numbers of pSMAD1/5/9-positive cells than wild-type (Fig. 1H). This suggests that at E5.5, BMP4 plays a major role in initiating BMP signaling activity in the mouse, and that this function of BMP4 is dose-dependent.

### Maternal *Bmp4* and *Smad4* are not required for development

Previous knockout studies of BMP signaling components did not report preimplantation phenotypes (Beppu et al., 2000; Mishina et al., 1995; Sirard et al., 1998; Winnier et al., 1995) However, other groups reported defects in preimplantation lineage specification using pathway inhibitors or microinjection of RNAi or mRNA for dominant-negative overexpression (Graham et al., 2014; Reyes de Mochel et al., 2015; Stuart et al., 2019) One way to reconcile these disparate findings is to invoke a model in which some components of the BMP pathway are maternally imparted to the oocyte and participate in preimplantation development to compensate for previously reported zygotic null mutations. To further investigate this possibility, we examined cell fate specification in embryos lacking both maternal (m) and zygotic (z) *Bmp4* or *Smad4* using the female germ line-expressed *Zp3*-*Cre* (De Vries et al., 2000) in combination with floxed alleles of either *Bmp4* or *Smad4* (see Supp. Fig. 3A for breeding scheme). RT-qPCR analysis confirmed the absence of detectable *Smad4* transcript in *Smad4* mz null embryos (Supp. Fig. 3B), as we have observed for many other loci deleted using this approach (Blij et al., 2012; Frum et al., 2013; Frum et al., 2018; Wicklow et al., 2014).

Remarkably, we were able to recover either *Bmp4* or *Smad4* mz null blastocysts at predicted rates, indicating no requirement for maternal *Bmp4* or *Smad4* on fertilization or embryo development. Moreover, both *Bmp4* and *Smad4* mz null embryos exhibited normal morphology, total cell number, and ratio of trophectoderm and ICM cells (Fig 2, Supp. Fig. 3C-E). In addition, the ICM marker *Oct4*, was detected at normal levels within the ICM, and CDH1 localization strongly suggested that the TE was properly polarized (Supp. Fig. 4). Finally, the expression of EPI and PrE cell fate markers at E3.75, E4.25, and E4.5 (Fig. 2, Supp. Fig. 3C-E) was also unaffected in either *Bmp4* or *Smad4* mz null embryos at these stages, consistent with normal ICM differentiation. Our observations support the conclusion that canonical BMP signaling does not play a major role in preimplantation development. In a parallel set of experiments, we allowed *Smad4* m null embryos to develop to term. Mice lacking m *Smad4* were born and developed apparently normally to 4 months old (6/6 mice, one litter). We conclude that maternal *Bmp4* and *Smad4* are dispensable for development and that neither zygotic gene plays a predominant role prior to implantation.

**Figure 2.**
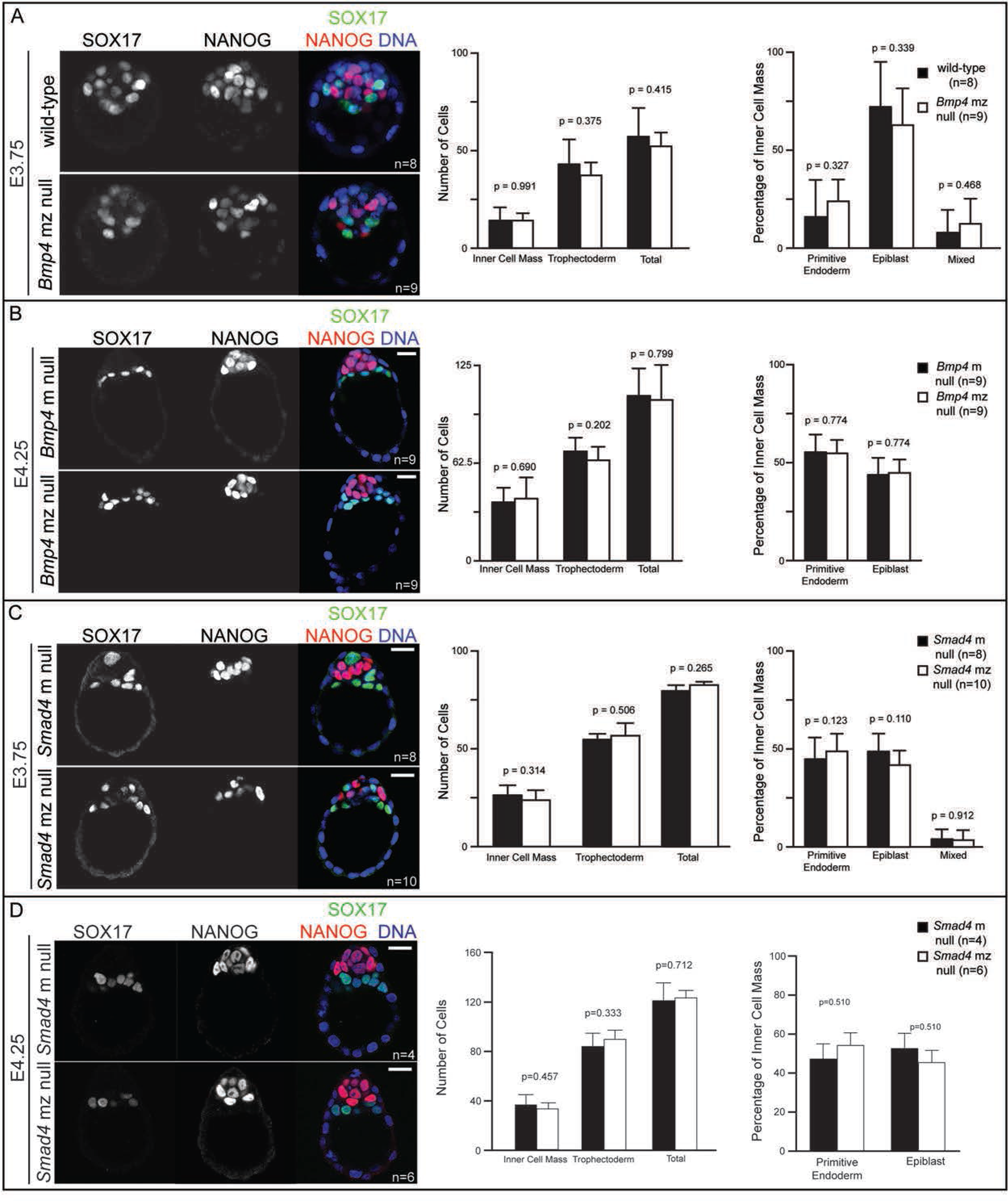
Maternal and zygotic *Smad4* and *Bmp4* are dispensable for blastocyst formation and preimplantation cell fate specification. A) Immunofluorescence for SOX17 and NANOG as respective markers of primitive endoderm (PrE) and epiblast (EPI) in flushed E3.75 wild-type CD-1 embryos and embryos lacking maternal and zygotic *Bmp4* (mz null). Quantification did not reveal any significant difference in cell number or cell fate between *Bmp4* mz null embryos and controls. “Mixed” indicates co-expression of SOX17 and NANOG. B) Immunofluorescence for SOX17 and NANOG in flushed E4.25 embryos lacking maternal *Bmp4* only (m null) and *Bmp4* mz null embryos. Quantification did not reveal any significant difference in cell number or cell fate between *Bmp4* mz null embryos and controls. “Mixed” indicates co-expression of SOX17 and NANOG. C) Immunofluorescence for SOX17 and NANOG in flushed E3.75 *Smad4* m null and *Smad4* mz null embryos. Quantification did not reveal any significant difference in cell number or cell fate between *Smad4* mz null embryos and controls. D) Immunofluorescence for SOX17 and NANOG in flushed E4.25 *Smad4* m null and *Smad4* mz null embryos. Quantification did not reveal any significant difference in cell number or cell fate between *Smad4* mz null embryos and controls. All pairwise comparisons were assessed by Student’s t-test. Scale bars represent 10 μm.

### *Smad4* is required for epiblast cavitation at E5.5 in a *Bmp4*-independent manner

Prior studies mainly focused on characterization of BMP signaling loss of function phenotypes at later stages (>E5.5) (Sirard et al., 1998; Winnier et al., 1995; Yang et al., 1998). However, we first observed BMP signaling activity in most embryos just after implantation at E4.75, prompting us to examine embryos lacking *Bmp4* or *Smad4* beginning at E4.75. At E4.75, *Smad4* null embryos were grossly morphologically normal (Fig. 3A). However, upon close examination, *Smad4* null embryos displayed a significant decrease in total cell number (Fig. 3B). We quantified the number of EPI, PrE, and TE cells in these embryos and discovered that the decreased cell number was most pronounced in EPI cells at this stage (Fig. 3B-C).

**Figure 3.**
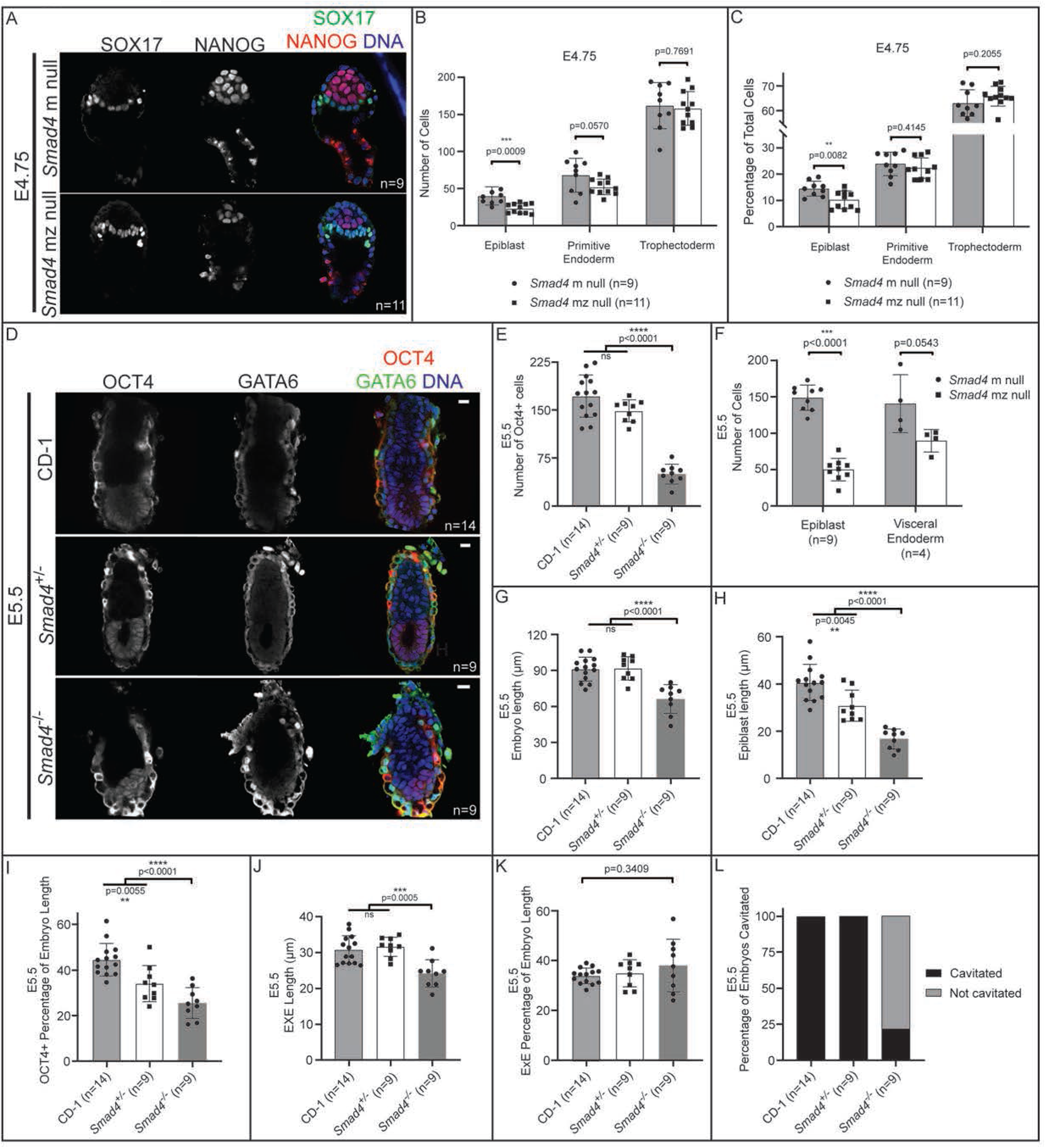
BMP-independent function of *Smad4* is required for post-implantation epiblast organization and maintenance. A) E4.75 *Smad4* mz null embryos stained by immunofluorescence for SOX17 and NANOG. B) Quantification of EPI, PrE, and TE cell numbers from embryos in A revealed a significant decrease in EPI cells in *Smad4* mz null embryos when compared to controls. C) Quantification of the EPI, PrE, and TE cells as a percentage of total cell number from embryos in A revealed a significant decrease in EPI percentage in *Smad4* mz null embryos. D) E5.5 *Smad4*^−/−^ embryos stained by immunofluorescence for OCT4 and GATA6 as markers of EPI and VE, respectively. *Smad4^−/−^* refers to combined *Smad4* z null and *Smad4* mz null embryos. E) Quantification of the number of OCT4+ cells in wild-type, *Smad4^+/−^,* and *Smad4^−/−^* embryos. F) Quantification of EPI and PrE cell numbers from *Smad4^+/−^* and *Smad4^−/−^* embryos at E5.5 revealed a specific, significant decrease in epiblast cell number in *Smad4* mz null embryos when compared to controls (p<0.05 by Student’s t-test). The difference in VE cell numbers was not significant (p>0.05). G) Quantification of the proximal-distal length wild-type, *Smad4^+/−^,* and *Smad4^−/−^* embryos at E5.5. H) Quantification of the proximal-distal length of the EPI of wild-type, *Smad4^+/−^,* and *Smad4^−/−^* embryos at E5.5. I) Quantification of the proximal-distal length of the EPI as a percentage of total length of wild-type, *Smad4^+/−^,* and *Smad4^−/−^*embryos at E5.5. J) Quantification of the proximal-distal length of the EXE of wild-type, *Smad4^+/−^,* and *Smad4^−/−^* embryos at E5.5. K) Quantification of the proximal-distal length of the EXE as a percentage of total length of wild-type, *Smad4^+/−^,* and *Smad4^−/−^* embryos at E5.5. L) Quantification of the proportion of *Smad4^+/−^* and *Smad4^−/−^*embryos with a proamniotic cavity at E5.5. Comparisons in B, C, F were assessed by Student’s t-test. Comparisons in E, G-K were assessed by analysis of variance (ANOVA) with Tukey’s post-hoc test.

By E5.5, *Smad4* null embryos were visibly reduced in size and all displayed disorganization in EPI, VE, and EXE compartments, as expected from studies performed at E6.5 (Sirard et al., 1998; Yang et al., 1998) (Fig. 3D). Strikingly, the epiblast was greatly reduced in cell number (Fig. 3D-F) relative to controls, and had not yet cavitated, as the proamniotic cavity was not present among most (>80%, n=7/9) *Smad4* null embryos examined (Fig. 3D, 3L). Notably, relative to extraembryonic lineages, the number of EPI cells was disproportionately decreased in *Smad4* null embryos since the epiblast length was decreased even when normalized to proximal-distal embryo length (Fig. 3G-I, Supp. Fig. 5A). By contrast, the size of the EXE was appropriately scaled to the reduced size of *Smad4* null embryos at E5.5 (Fig. 3J-K). This suggests that *Smad4* is not only required for general embryonic growth, but also specifically required for epiblast growth relative to total embryo size. Notably, these phenotypes were not observed in E5.5 *Bmp4-*null embryos, which did not differ from wild-type in morphology, embryo size, or EPI or PrE cell number (Supp. Fig. 5B-E), suggesting that alternative signaling pathways feed into SMAD4-regulation of epiblast growth and morphogenesis at this stage.

### Epiblast cavitation requires SMAD4-dependent inhibition of FGF/ERK signaling

Having discovered that *Smad4* is required for epiblast cavitation and growth at E5.5, we began to investigate the mechanism underlying this role. We were struck by the observation that the EXE appeared disproportionately large, relative to the size of the EPI in E5.5 *Smad4* null embryos. To confirm the identity of the EXE cells, we examined markers of EXE, including phosphorylated ERK (pERK), which is elevated within the EXE (Corson et al., 2003). In *Smad4* null embryos, we observed pERK throughout putative EXE, consistent with their identity as EXE cells. However, we observed dramatically elevated levels of pERK within this region (Fig. 4A). Since pERK in the EXE is known to be dependent on signaling by the Fibroblast Growth Factor (FGF) pathway (Corson et al., 2003), we therefore hypothesized that increased pERK could be due to elevated FGF signaling in *Smad4* null embryos.

**Figure 4.**
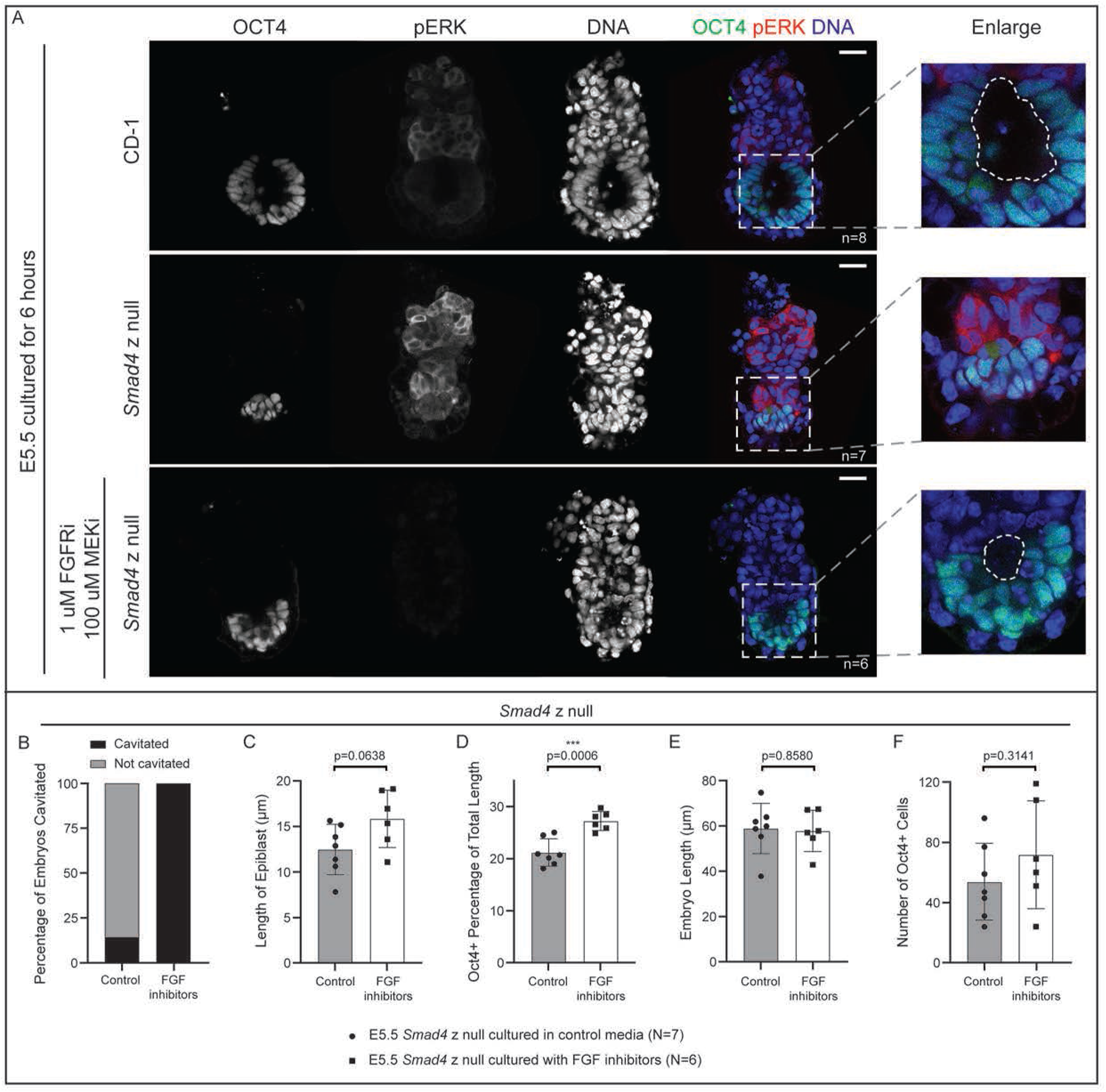
Inhibition of FGF signaling partially rescues epiblast cavitation in E5.5 *Smad4* null embryos. A) Wild-type and *Smad4* z null embryos collected at E5.5 and cultured for 6 hours after dissection in control media or media containing FGFR/MEK inhibitors (see Methods), then stained by immunofluorescence for OCT4 and phosphorylated ERK (pERK). Dashed line in enlargement denotes the proamniotic cavity. B) Quantification of the proportion of treated and untreated *Smad4^−/−^* embryos with a proamniotic cavity at E5.5. C) Quantification of proximal-distal length of the EPI in treated and untreated E5.5 *Smad4^−/−^* embryos. D) Quantification of proximal-distal length of the EPI as a proportion of total length in treated and untreated E5.5 *Smad4^−/−^* embryos. E) Quantification of proximal-distal length in treated and untreated E5.5 *Smad4^−/−^* embryos. F) Quantification of OCT4-positive cell number in treated and untreated E5.5 *Smad4^−/−^* embryos.

To test this hypothesis, we used a previously-published protocol to inhibit FGF signaling in embryos (Yamanaka et al., 2010), which effectively eliminated pERK in control embryos (Supp. 6A). Notably, inhibition of FGF signaling partially rescued epiblast defects in E5.5 *Smad4* null embryos (Fig. 4A). We observed a significant increase in epiblast scaling, as well as increased rates of cavitation (Fig. 4B-D). However, FGF inhibitor treatment did not rescue the overall growth restriction of *Smad4*-null embryos (Fig. 4E), nor was the number of OCT4-positive cells restored (Fig. 4F). These observations are consistent with SMAD4 repressing FGF signaling in the EXE during early post-implantation stages as a critical regulator of epiblast morphogenesis. Our observations also suggest that additional pathways regulate embryo growth downstream of SMAD4.

To determine whether elevated FGF signaling is sufficient to antagonize cavitation, we treated embryos with exogenous FGF4. In wild-type E5.5 embryos treated with exogenous 1 µg/mL FGF4, we observed elevated levels of pERK within the EXE and ectoplacental cone (Supp. Fig. 6A). However, we observed no impact on epiblast cavitation or epiblast size following this treatment (Supp. Fig. 6A-D). Altogether, these data suggest that pERK/FGF signaling antagonizes EPI cavitation, but upregulation of pERK alone is insufficient to induce cavitation defects in wild-type embryos, at least under the conditions tested here.

## Discussion

Despite being a highly conserved developmental signaling pathway, the role of BMP signaling in regulating peri-implantation mammalian development has not been fully elucidated. In part, this may be due to the technical challenges surrounding studying mammalian embryos at pre- and peri-implantation stages of development. Our study undertook a detailed examination of molecular markers of BMP signaling activity and the phenotypes of maternal-zygotic genetic knockout models of BMP signaling to identify the earliest role of BMP signaling in development. Although we failed to detect a role for BMP signaling in preimplantation embryos, we identified a novel role for *Smad4* in epiblast organization and cavitation at early post-implantation stages.

We report a novel requirement for *Smad4* in post-implantation development. Previous descriptions of *Smad4*-null embryos report growth restriction and disorganized VE beginning at E5.5, with embryonic lethality by E8.5 (Chu et al., 2004; Sirard et al., 1998; Yang et al., 1998). Our results are consistent with these findings, but also uncover a previously unappreciated defect specific to the epiblast. The proportion of EPI cells are decreased starting at E4.75, suggesting that in the absence of *Smad4* the number of EPI cells does not scale with embryo growth. The structure of the epiblast is also disrupted in these embryos, as *Smad4*-null embryos had largely disorganized epiblasts which failed to form a proamniotic cavity. It is unlikely that this defect is cell-autonomous, as embryos with an epiblast-specific knockout of *Smad4* do not display an observable defect until after gastrulation (Chu et al., 2004). Rather, it is likely that *Smad4* is required in the visceral endoderm to produce a signal that promotes EPI development non-cell-autonomously (Fig. 5A). This model is supported by the finding that VE-specific *Smad4*-knockout embryos fail to gastrulate properly and more closely resemble whole-body *Smad4-*null embryos (Li et al., 2010).

**Figure 5.**
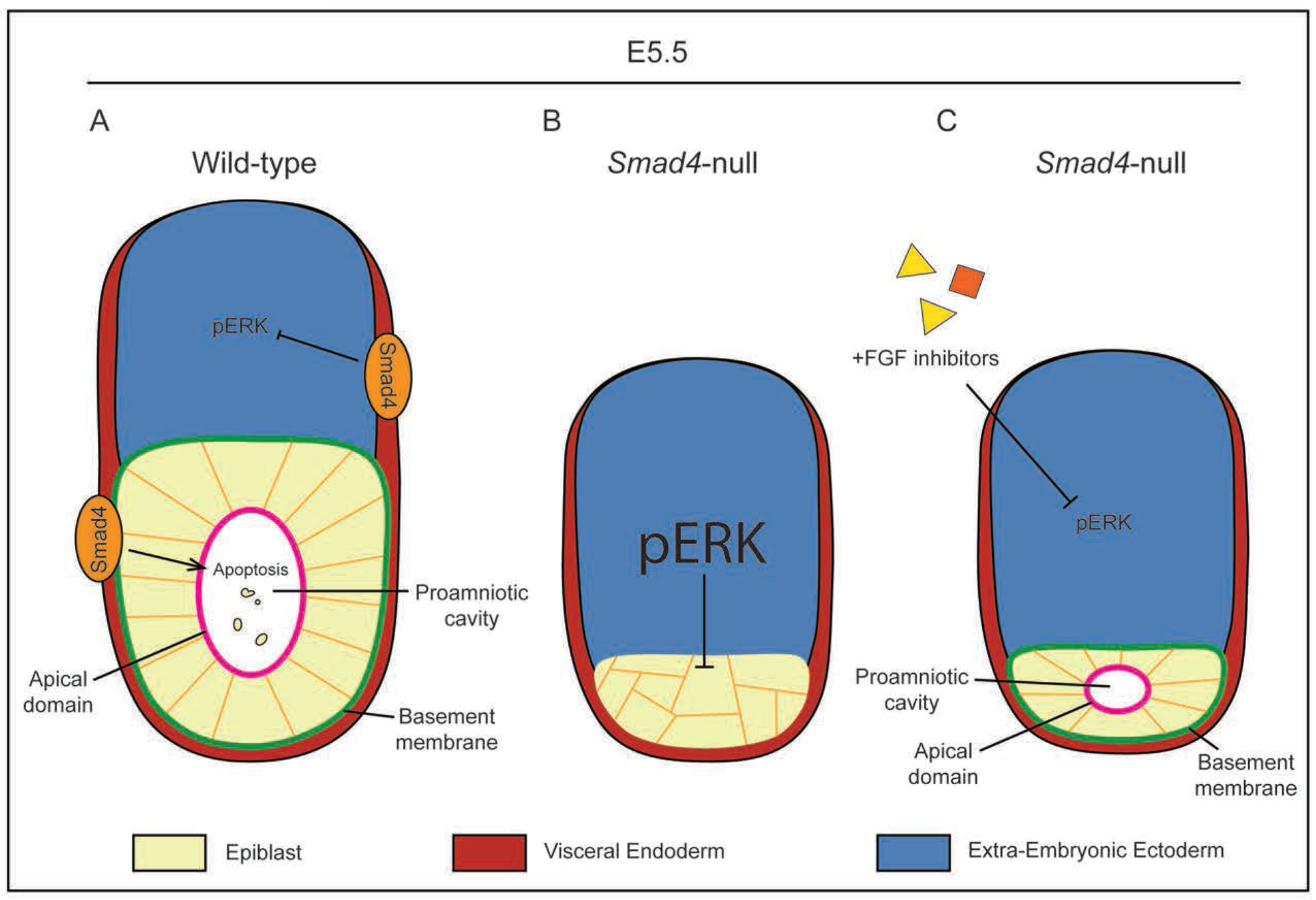
FGF inhibition rescues rosette formation but not embryo growth in *Smad4*-null embryos. A) In wild-type embryos, SMAD4 in the visceral endoderm promotes lumen formation in the epiblast by regulating a pro-apoptotic signal. Separately, SMAD4 also inhibits ERK phosphorylation in the EXE, which allows for polarization and rosette formation in the epiblast. B) In *Smad4*-null embryos, pERK is upregulated, causing an ectopic increase in pERK and preventing epiblast polarization. The pro-apoptotic signal is also lost. The combination of these two factors lead to epiblast disorganization and a failure to cavitate. C) Treatment with FGF inhibitors prevents upregulation of pERK in *Smad4*-null embryos. Repressed pERK levels allow epiblast polarization to proceed, resulting in a small proamniotic cavity.

Our observations are consistent with prior observation that BMP signaling is necessary and sufficient for cavitation of embryoid bodies (Coucouvanis and Martin, 1999). In this context, BMP2 and BMP4 may be functionally redundant, and could explain why we observed normal cavitation in E5.5 *Bmp4*-null embryos. Alternatively, SMAD4 could regulate epiblast morphogenesis through another TGF-β pathway. Knockout of *Nodal* has been shown to decrease embryo size and expression of *Oct4* mRNA at early post-implantation stages (Brennan J. et al., 2001; Mesnard et al., 2006), but *Nodal-*null embryos cavitate normally and expression of OCT4 protein is apparently unaffected at E5.5 (Senft et al., 2019). As the cavitation defect and loss of OCT4-positive cells are more severe in *Smad4-*null embryos than either *Bmp4-* or *Nodal-*null models alone, it suggests that *Smad4* may regulate epiblast morphogenesis through a combination of TGFβ pathways.

Our data also suggests that SMAD promotes cavitation by attenuating pERK levels. Inhibition of FGF/MAPK signaling rescued cavitation in *Smad4-*null embryos, even though the epiblast remained disproportionately small. This observation allows us to propose that cavitation is not dependent on epiblast size, but rather the embryo signaling environment. Our data suggest that ectopic upregulation of pERK resulting from loss of *Smad4* is detrimental to epiblast cavitation; however, the mechanisms of how these two pathways interact with one another and regulate cavitation remain unclear. SMAD4 could regulate several processes associated with epiblast cavitation, including epiblast cell apoptosis or epiblast cell polarization and lumenogenesis (Bedzhov and Zernicka-Goetz, 2014; Coucouvanis and Martin, 1995; Coucouvanis and Martin, 1999; Halimi et al., 2022) (Fig 5B-C). Alternatively, SMAD4 could promote epiblast maturation, defined here as the transition from a naïve to primed state which normally occurs in epiblast cells between preimplantation and post-implantation stages (Boroviak et al., 2015; Nichols and Smith, 2009). Epiblast maturation has been shown to be critical in formation of proamniotic cavity (Carbognin et al., 2023; Shahbazi et al., 2017). Other mechanisms are also proposed to play a role in embryonic cavitation such as apical domain maintenance (Meng et al., 2017), tight junction formation (Chan et al., 2019), cellular adhesion through ECM interactions (Liang et al., 2005; Sakai et al., 2003), and establishment of an osmotic gradient (Dumortier et al., 2019). SMAD4 and pERK may regulate any or a combination of these factors. We note that experimental elevation of pERK was not sufficient to inhibit cavitation in wild-type embryos, at least under the conditions tested here. This observation could suggest that pERK regulates expression of components of the cavitation process that are not themselves rate-limiting. Further studies will be needed to interrogate the interaction of these pathways further, to identify the precise regulatory mechanisms governing post-implantation epiblast morphogenesis downstream of SMAD4 and pERK.

## Materials and Methods

**Table 1.**
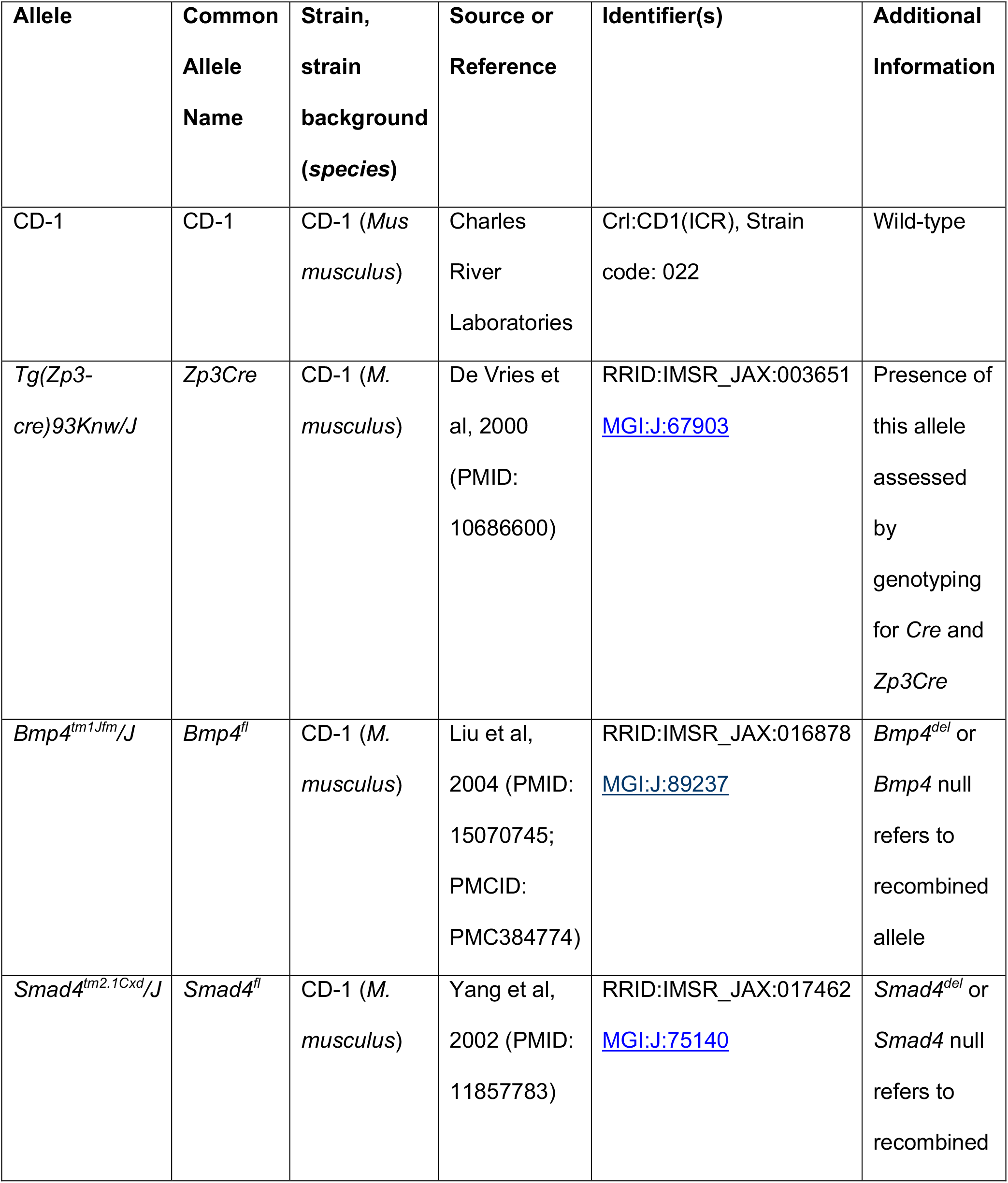
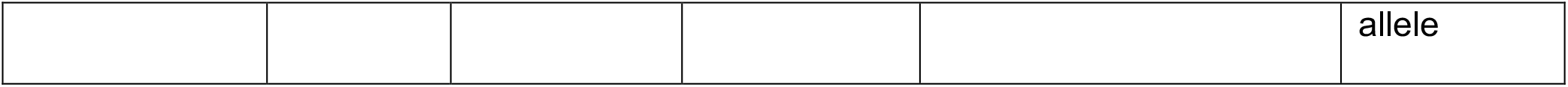
Animal Resources Table.

**Table 2.**
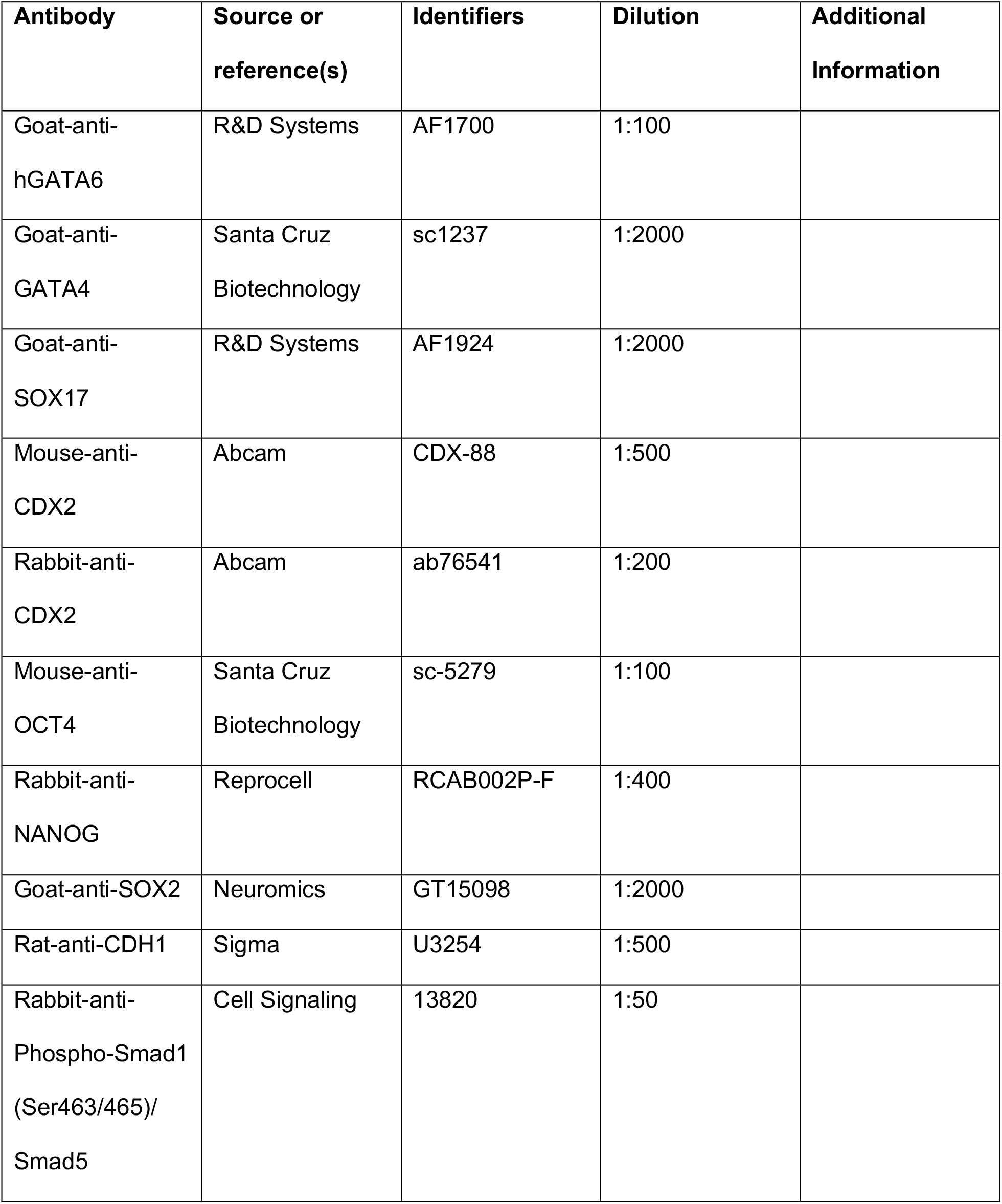
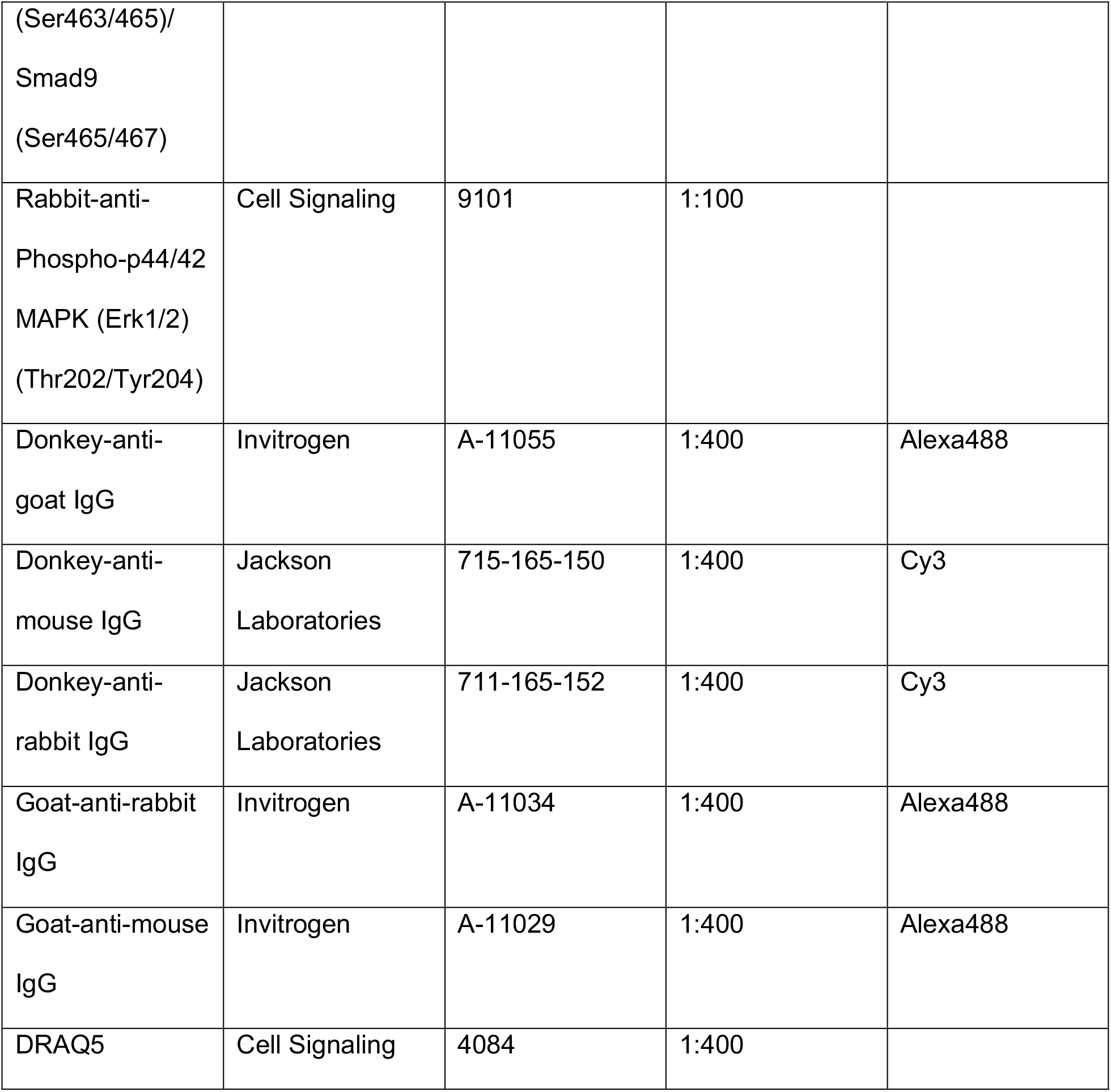
Antibody Table.

**Table 3.**
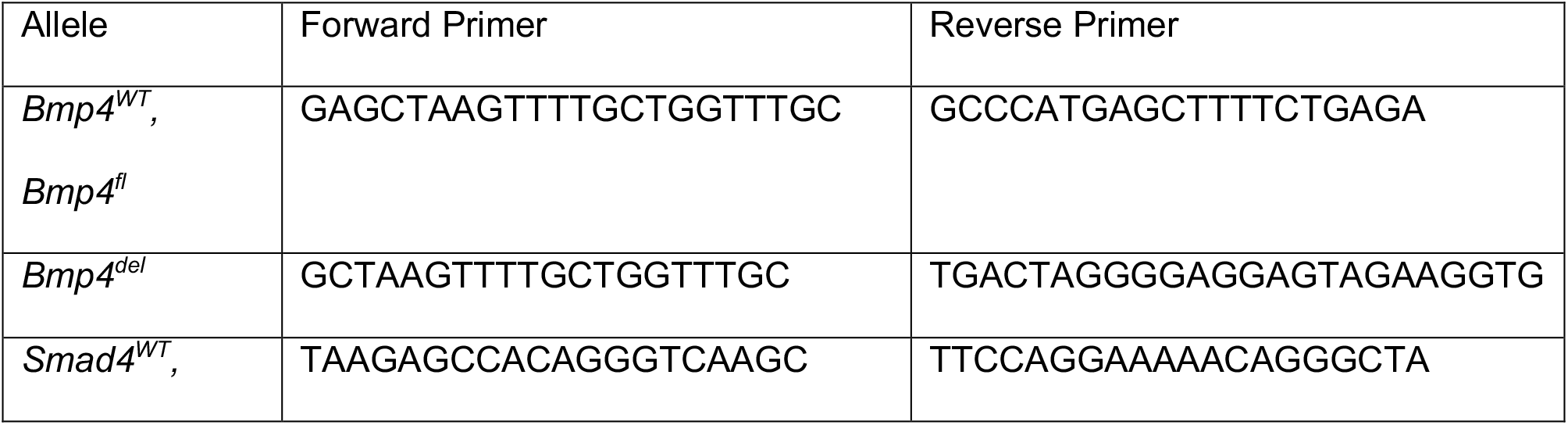
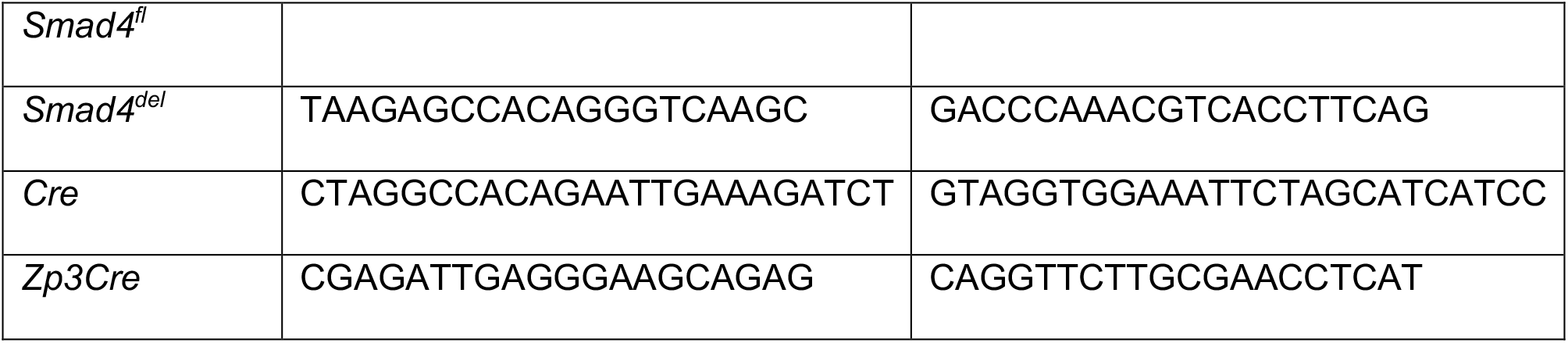
Allele-specific primers for PCR genotyping.

### scRNA-seq Analysis

Single-cell RNA-seq data generated by Nowotschin *et. al*. was used to analyze the expression of TGFβ genes in mouse E3.5, E4.5, E5.5, and E6.5 blastocysts (Nowotschin et al., 2019) (Nowotschin et al., 2019). The analysis was completed using R v4.1.0 with tools from Seurat v4.3.0 (R-core Team 2021) (Hao et al., 2021). We normalized the UMI counts using SCTransform and cells were visualized in 2D space using UMAP performed on the first 30 principal components (Choudhary and Satija, 2022; Hafemeister and Satija, 2019). After excluding TGFβ genes expressed in <10 cells, we used Seurat’s FindAllMarkers function with the Wilcoxon rank-sum test to identify TGFβ genes enriched in each cell type versus all other cells. The p-values were corrected for multiple comparisons using the Bonferroni method. Genes with p-adj<0.01 and average log_2_ fold change<0.25 were considered cluster enriched. Heatmaps were generated using the pheatmap (v 1.0.12) after averaging the normalized expression for each gene in each cell type.

### Mouse Strains and Genotyping

All animal research was conducted in accordance with the guidelines of the Michigan State University Institutional Animal Care and Use Committee. Wild type embryos were derived from CD-1 mice (Charles River). The following alleles were used in this study and maintained in a CD-1 background: *Bmp4^tm1Jfm^/J* (Liu et al., 2004); *Smad4^tm2.1Cxd^/J* (Yang et l., 2002); *Tg(Zp3-cre)93Knw* (de Vries et al., 2000). Null alleles were generated by breeding dams carrying homozygous floxed alleles and the *Zp3Cre* allele to CD-1 males. Mouse genotypes were determined by PCR using genomic DNA extracted using the REDExtract-N-Amp kit (Sigma XNAT) according to the manufacturer’s protocol. Embryo genomic DNA was extracted using the same kit scaled to 10 µL total volume. Genomic extracts (1–2 µL) were then subjected to PCR using allele-specific primers (see Table 3).

### Embryo Collection and Culture

Mice were maintained on a 12 hr light/dark cycle. Preimplantation (E2.5-E4.5) embryos were collected by flushing the oviduct or uterus with M2 medium (Sigma M7167). Post-implantation (E4.75-E6.5) embryos were collected by dissecting the embryos from the decidua in ice-cold PBS containing 1% FBS (HyClone SH30396.02) or Bovine Serum Albumin (BSA, Sigma A7888). During embryo collection, dissected embryos were held in warm M2 media. For embryo culture, KSOM medium (Millipore MR-121-D) was equilibrated overnight prior to embryo collection. Where indicated, the following were included in the culture medium: 1 µM or 0.25 µM LDN-193189 in DMSO (Stemgent 04-0074-02); 1 µg/mL recombinant FGF4 in PBS with 0.1% BSA (R&D 235-F4); 1 µg/mL heparin (Sigma H3149); 100 ng/mL recombinant BMP4 in 4 mM HCl (R&D 314-BP); 1 µM PD173074 in DMSO (Selleckchem S1264); 5 µM PD0325901 in DMSO (Stemgent 04-0006); or 0.2% DMSO (New England BioLabs B0515A) as control. Embryos were cultured at 37°C in a 5% CO_2_ incubator under light mineral oil (Millipore ES-005-C).

### Real-time PCR of Oocytes

*Smad4* expression levels in oocytes were assessed by real-time PCR as previously described (Blij et al, 2012). *Smad4* levels were assessed in oocytes from three wild-type and three *Smad4* maternal null females. Oocytes collected from each female were pooled for mRNA extraction and cDNA synthesis. RT-PCR was performed in quadruplicate technical replicates for each cDNA sample. Primers were (5′-3′): *Actb*, CTGAACCCTAAGGCCAACC and CCAGAGGCATACAGGGACAG; *Smad4* (wild-type allele) CGCGGTCTTTGTACAGAGTTA and ACACTGCCGCAGATCAAAG; *Smad4* (deleted allele), CACAGGACAGAAGCGATTGA and CCAAACGTCACCTTCACCTT.

### Immunofluorescence and Confocal Microscopy

Preimplantation embryos (E2.5-E4.75) were fixed with 4% formaldehyde (Polysciences 04018) for 10 min, permeabilized with 0.5% Triton X-100 (Sigma Aldrich X100) for 30 min, and then blocked with blocking solution (10% Fetal Bovine Serum (HyClone SH30396.02), 0.1% Triton X-100) overnight at 4°C. Embryos were incubated with primary antibody overnight at 4°C. The next day, embryos were washed in blocking solution for 30 min, incubated in secondary antibody diluted in blocking solution for 1 hr, washed in blocking solution for 30 min, then stained with nuclear stain diluted in block for 10 min or overnight.

Post-implantation embryos (E5.0-E5.75) were fixed with 4% formaldehyde for 1 hr, washed 3 times in 0.1% Tween-20 (Sigma Aldrich P9416), permeabilized for 4 hrs in 0.5% Triton X-100, and then blocked with blocking solution (3% BSA (Sigma Aldrich A7888); 0.3% Triton X-100 in PBS) overnight at 4°C. Embryos were incubated with primary antibody overnight at 4°C. The next day, embryos were washed three times in 0.1% Tween-20 for 5 min, then incubated in secondary antibody diluted in blocking solution overnight. The following day embryos were washed three times in 0.1% Tween-20 for 5 min, then stained with nuclear stain diluted in block for 10 min or overnight.

All embryos (preimplantation or post-implantation) which used antibodies against pSMAD1/5/9 were fixed with 4% formaldehyde for 1 hr, methanol dehydration-rehydration series (25%, 50%, 75%, 100%) for 5 min each, washed three times in freshly-made 1% Triton X-100 for 10 min, washed 20 min in ice-cold acetone at −20°C, washed three times in freshly-made 1% Triton X-100 for 10 min, then then blocked with blocking solution (10% Fetal Bovine Serum, 0.1% Triton X-100 in PBS) overnight at 4°C. Embryos were incubated with primary antibody overnight at 4°C. The next day, embryos were washed three times in freshly-made 0.1% Triton X-100 for 10 min, incubated in secondary antibody diluted in blocking solution for 2 hrs, washed three times in freshly-made 0.1% Triton X-100 for 10 min, then stained with nuclear stain diluted in blocking solution for 10 min or overnight.

All embryos (preimplantation or post-implantation) which used antibodies against pERK were fixed with 4% formaldehyde for 1 hr, washed three times for 5 min in PBS, washed 20 min in ice-cold methanol at −20°C, permeabilized 30 min in 0.1% Tween-20, then blocked with blocking solution (3% Bovine Serum Albumin; 0.3% Triton X-100 in PBS) overnight at 4°C. Embryos were incubated with primary antibody overnight at 4°C. The next day, embryos were washed three times in PBS for 5 min, incubated in secondary antibody diluted in blocking solution for 2 hrs, washed three times in PBS for 5 min, then stained with nuclear stain diluted in block for 10 min or overnight. All solutions contained HALT protease inhibitor (Thermo Scientific 78430) and PhosSTOP phosphatase inhibitor (Roche 04906837001) diluted 1:500.

Antibodies used are listed in Table 2. Embryos were imaged using an Olympus FluoView FV1000 Confocal Laser Scanning Microscope system with 60X PlanApoN oil (NA 1.42) objective. For each embryo, z-stacks were collected, with 5 µm intervals between optical sections. All embryos were imaged prior to knowledge of their genotypes.

### Embryo Analysis

For each embryo, z-stacks were analyzed using Fiji (ImageJ), which enabled the labeling, based on DNA stain, of all individual cell nuclei. Using this label to identify individual cells, each cell in each embryo was then assigned to relevant phenotypic categories, without knowledge of embryo genotype. Phenotypic categories included marker expression (e.g., OCT4 positive or negative) and marker localization (e.g., pSMAD1/5/9 nuclear, absent, or unlocalized). Statistical analysis was performed using GraphPad Prism (v. 9.5.1). Figure images were assembled using Adobe Illustrator.

## Acknowledgements

We thank the lab of Dr. David Arnosti for their generous loan of laboratory equipment. We also thank Barbara Makela and Ella Markley for technical support.

## Diversity and Inclusion Statement

The authors wholeheartedly support all efforts to increase the inclusion of scientists from underrepresented backgrounds (including, but not limited to, women, LGBTQIA+, people of color, Indigenous people, neurodivergent people, people with disabilities, and people from disadvantaged backgrounds) in developmental biology and related careers. This manuscript reflects the efforts of authors who identify as members of several of these groups. The authors believe that diverse perspectives are essential for scientific excellence and innovation, yet acknowledge the continued existence of systemic barriers to success for scientists of underrepresented and marginalized communities. We support the development of initiatives to address disparities and biases in scientific publishing and encourage further efforts to implement inclusive practices.

