## Supplemental for "*Smad4* is essential for epiblast scaling and morphogenesis after implantation, but nonessential prior to implantation in the mouse"

#### Supplemental Figure Legends

**Supplemental Figure 1. SMAD1/5/9 phosphorylation in post-implantation embryos can be modulated.**

A) Immunofluorescence for GATA6 and SMAD1/5/9 phosphorylation (pSMAD1/5/9) in wild-type E4.25, E5.0, and E5.75 embryos. B) Immunofluorescence for GATA6 and pSMAD1/5/9 in wild-type E5.5 embryos after 6 hours of culture in control media or BMP inhibitor treatments. Note the lack of pSMAD1/5/9 with 0.25 μM LDN-193189 treatment. 1 μM LDN-193189 resulted in severe toxicity to the treated embryos. Graph represents quantification of the proportion of embryos displaying unique SMAD1/5/9 phosphorylation patterns. C) Immunofluorescence for GATA6 and pSMAD1/5/9 in wild-type E5.5 embryos after 6 hours of culture in indicated concentrations of exogenous BMP4 treatment. pSMAD1/5/9 showed a dose-dependent increase as BMP4 concentration increased. Graph represents quantification of proportion of embryos displaying unique SMAD1/5/9 phosphorylation patterns. “Proximal VE only” and “Proximal VE and DVE” refer to pSMAD1/5/9 signal in some cells in those tissues, while “Entire VE” refers to pSMAD1/5/9 signal in every observed VE cell. pSMAD1/5/9 signal in proximal VE is indicated by white arrowheads. pSMAD1/5/9 signal in DVE is indicated by yellow arrowheads. pSMAD1/5/9 signal in proximal EPI is indicated by red arrowhead.

**Supplemental Figure 2. BMP pathway genes are upregulated after implantation.**

A) Single-cell RNA-seq data generated by Nowotschin *et al*. reclustered using Seurat and identified by cell type. B) BMP ligand genes enriched in identified cell types and stages. Genes with p-adj<0.01 and average $\log_2$ fold change <0.25 were considered cluster enriched (*).

**Supplemental Figure 3. Maternal and zygotic *Smad4* and *Bmp4* are dispensable for blastocyst formation and preimplantation cell fate specification.**

A) Breeding scheme for the generation of *Smad4* maternal-zygotic (mz) null embryos using the *Zp3Cre* allele described in de Vries *et al*, 2000. The same strategy was applied to generate *Bmp4* mz null embryos. B) Level of *Smad4* mRNA detected in wild-type and *Zp3Cre*; *Smad4*$^{fl/fl}$ oocytes by qPCR, normalized to *Actin* mRNA level. C) Immunofluorescence for SOX17 and NANOG in dissected E4.5 *Smad4* m null and *Smad4* mz null embryos. D) Quantification of the number of cells occupying the inner cell mass or trophectoderm and total cell number in embryos from C. E) Quantification of the ratio of EPI and PrE cells in the inner cell mass of embryos from C. Pairwise comparisons were evaluated by Student's t-test.

**Supplemental Figure 4. Maternal and zygotic *Smad4* are not required for OCT4 expression in preimplantation mouse embryos.**

A) Immunofluorescence for OCT4 and CDH1 in E2.75 *Smad4* m null and *Smad4* mz null embryos. B) Quantification of the number of OCT4-positive cells as a percentage of total cells in embryos from A. C) Immunofluorescence for OCT4 and CDH1 in E3.75 *Smad4* m null and *Smad4* mz null embryos. D) Quantification of the number of OCT4-positive cells as a percentage of total cells in embryos from B. E) Immunofluorescence for OCT4 and CDH1 in E4.25 *Smad4* m null and *Smad4* mz null embryos. F) Quantification of the number of OCT4-positive cells as a percentage of total cells in embryos from E. All comparisons were assessed by analysis of variance (ANOVA).

**Supplemental Figure 5. *Bmp4* is dispensable for embryo growth and organization at E5.5.**

A) Correlation between the number of OCT4+ cells and proximal-distal length of the EPI in wild-type, *Smad4*$^{+/-}$, and *Smad4*$^{-/-}$ embryos at E5.5 from Fig. 3 and Fig. 4. Correlation assessed by Pearson's coefficient. B) Immunofluorescence for GATA6 and CDX2 as respective markers of visceral endoderm and extra-embryonic ectoderm in E5.5 wild-type, *Bmp4*$^{+/-}$, and *Bmp4*$^{-/-}$ embryos. C) Quantification of the proximal-distal length of embryos from B. D) Quantification of the proximal-distal length of the EPI in embryos from B. E) Quantification of the proximal-distal length of the EPI as a percentage of total length in embryos from B.

**Supplemental Figure 6. ERK phosphorylation is responsive to modulation of FGF signaling.**

A) Wild-type embryos collected at E5.5 and cultured for 6 hours in control media, 1 ug/mL FGF4 + heparin, or media containing FGFR/MEK inhibitors (see Methods), then stained by immunofluorescence for OCT4 and phosphorylated ERK (pERK). Note decreased pERK in inhibitor-treated embryos and increased pERK in FGF4-treated embryos. B) Quantification of proximal-distal length of all embryos from Figure 4A and Supplemental Figure 6A. C) Quantification of proximal-distal length of the EPI in all embryos from Figure 4A and Supplemental Figure 6A. D) Quantification of proximal-distal length of the EPI as a percentage of total length in all embryos from Figure 4A and Supplemental Figure 6A. All comparisons were assessed by analysis of variance (ANOVA) with Tukey's post-hoc test.

### Supplemental Figure 1

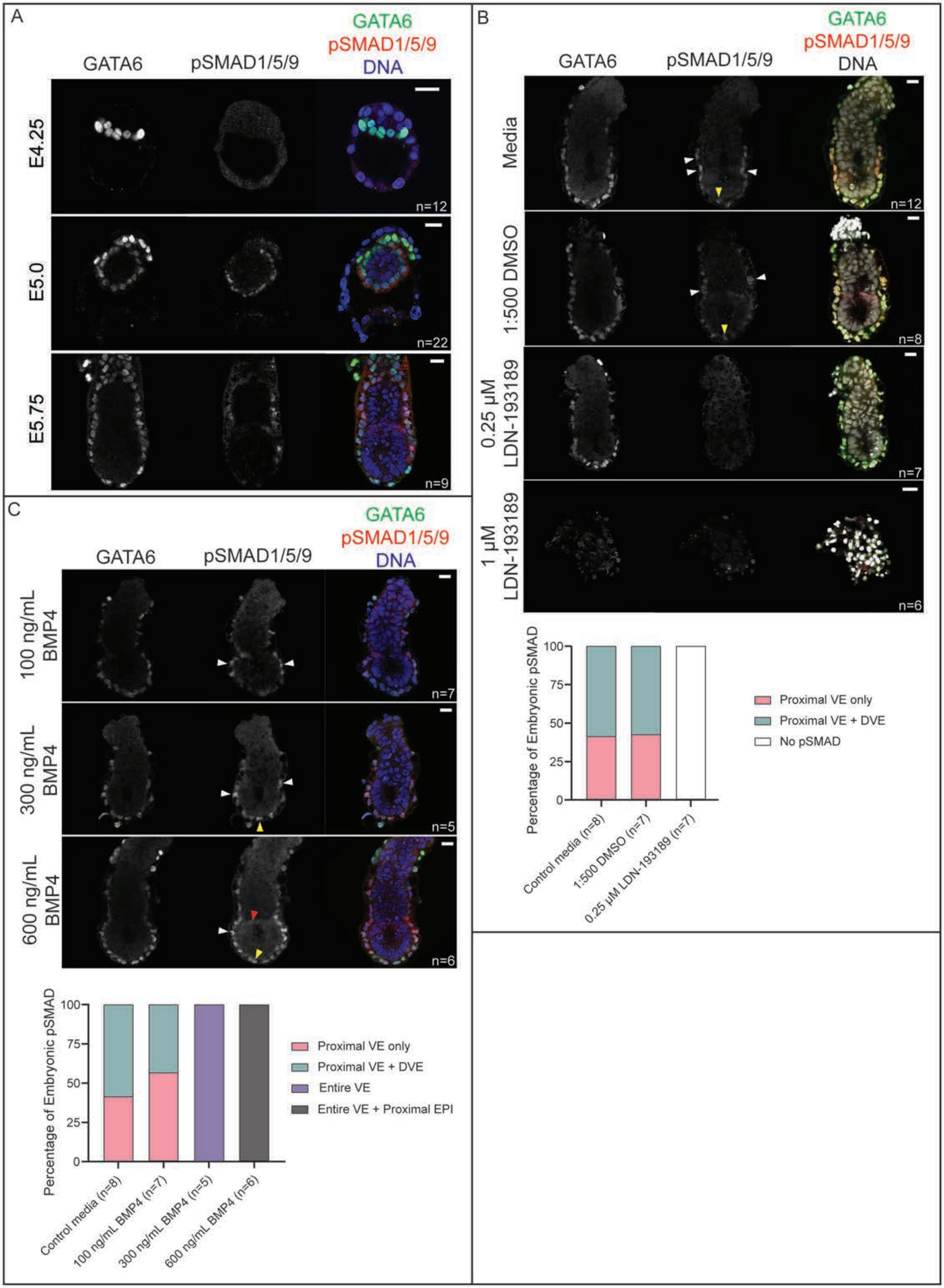

### Supplemental Figure 2

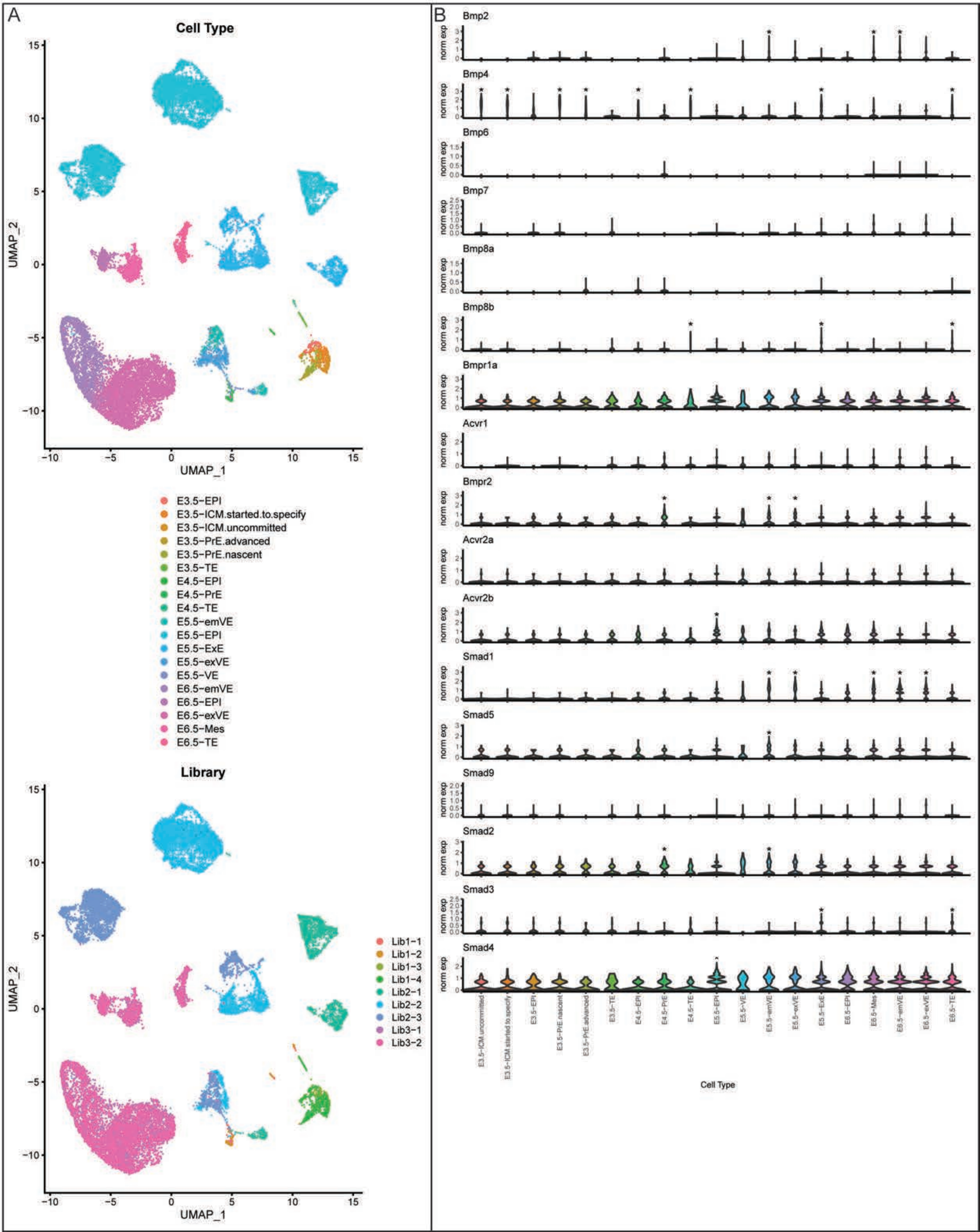

### Supplemental Figure 3

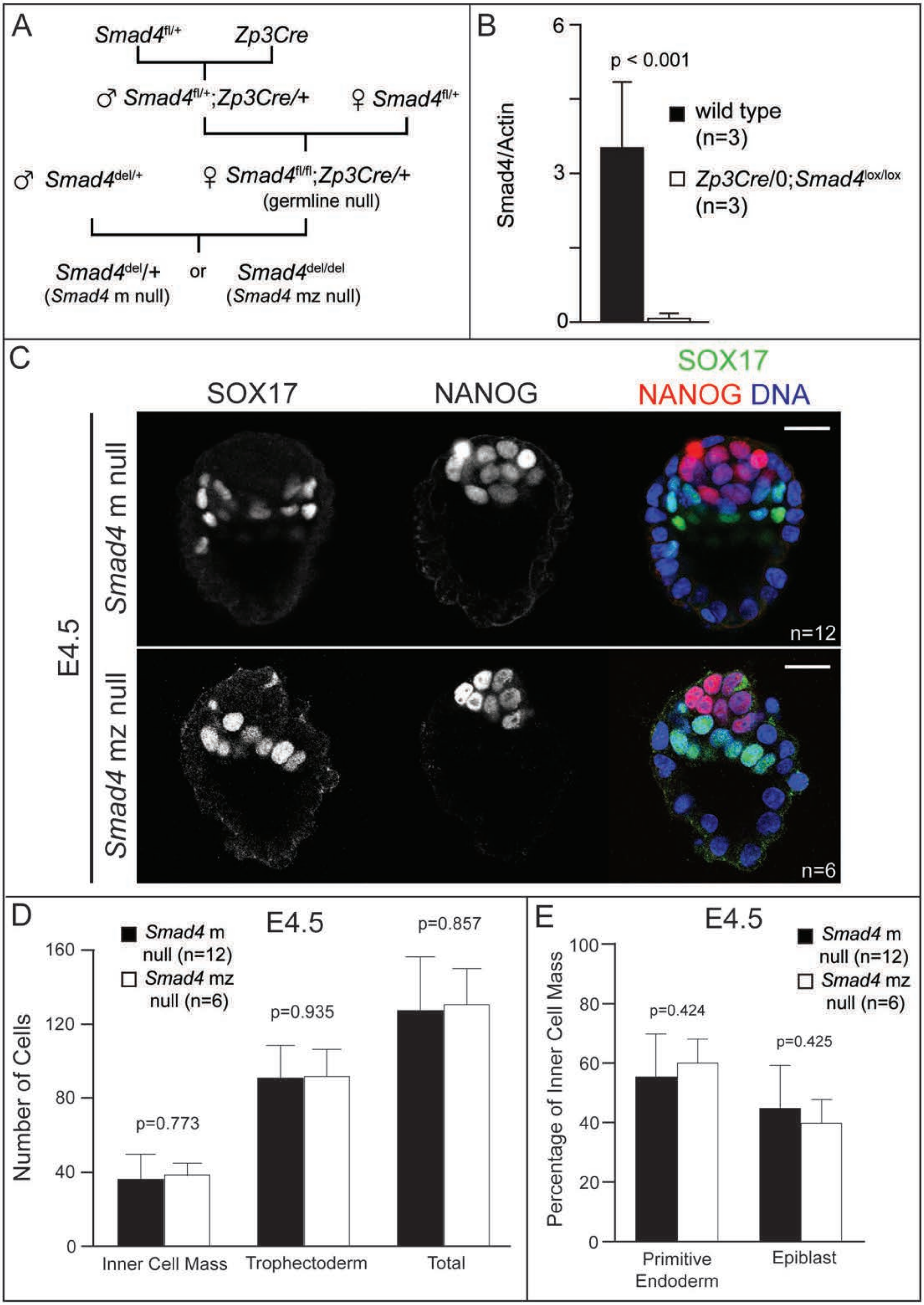

### Supplemental Figure 4

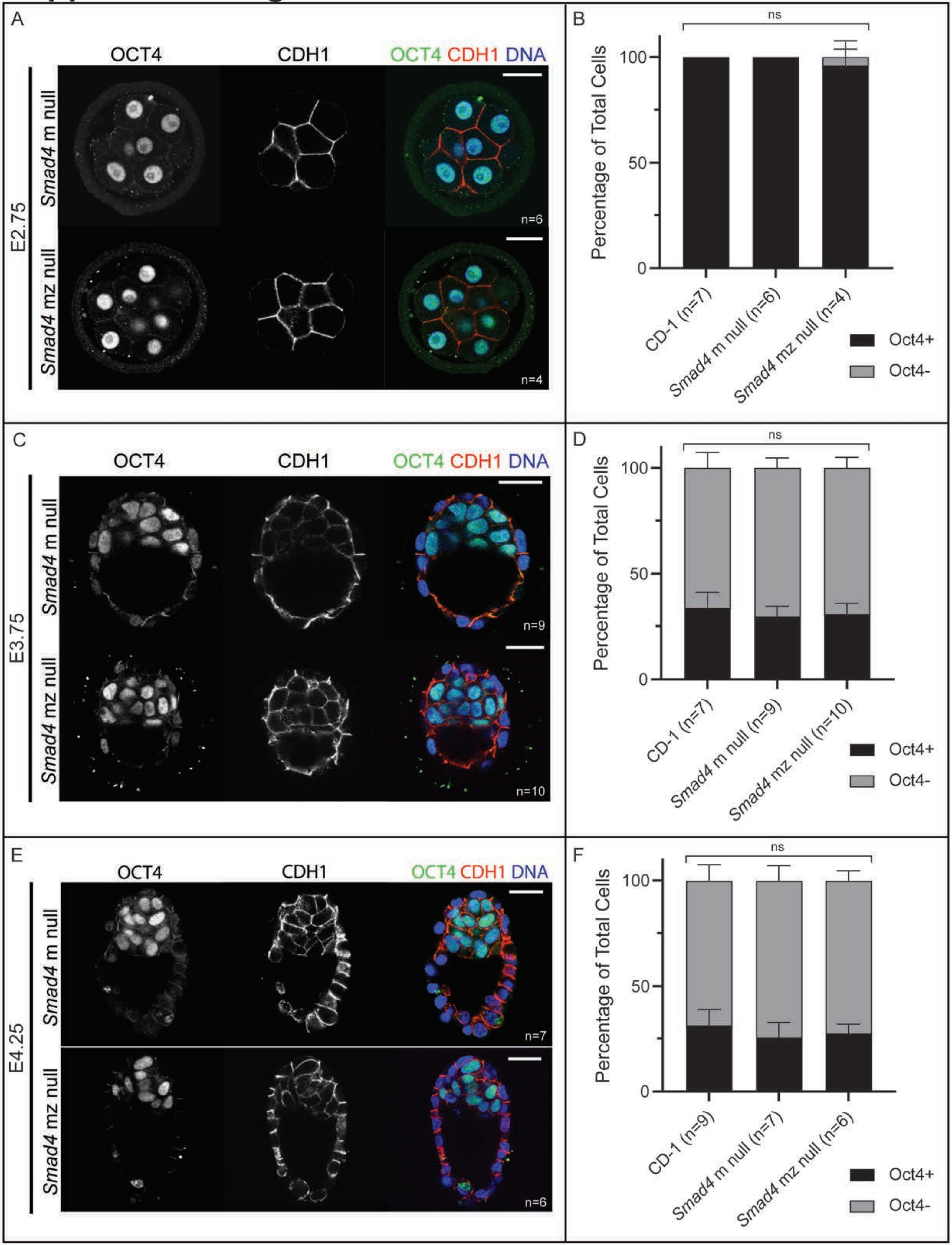

### Supplemental Figure 5

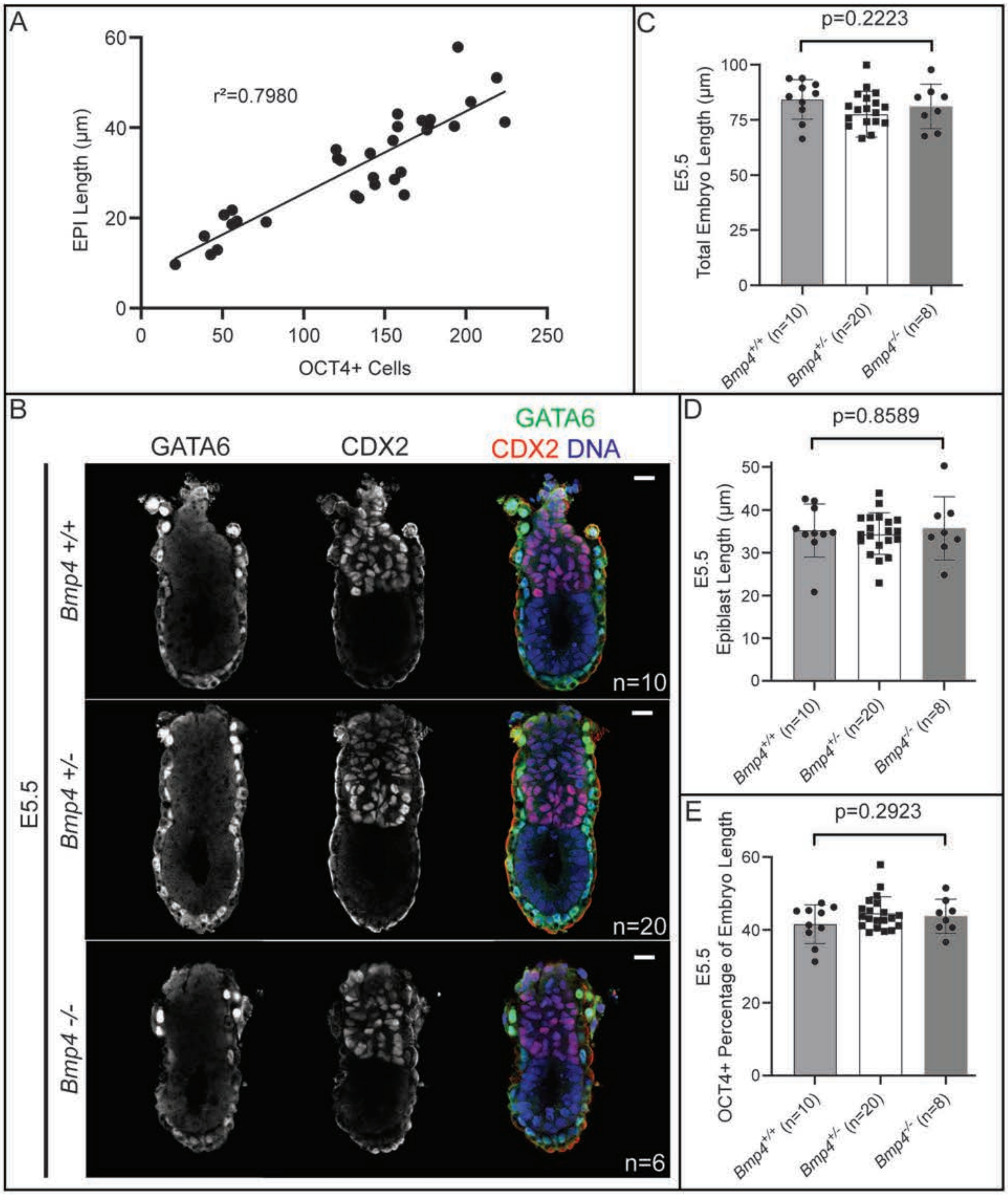

### Supplemental Figure 6

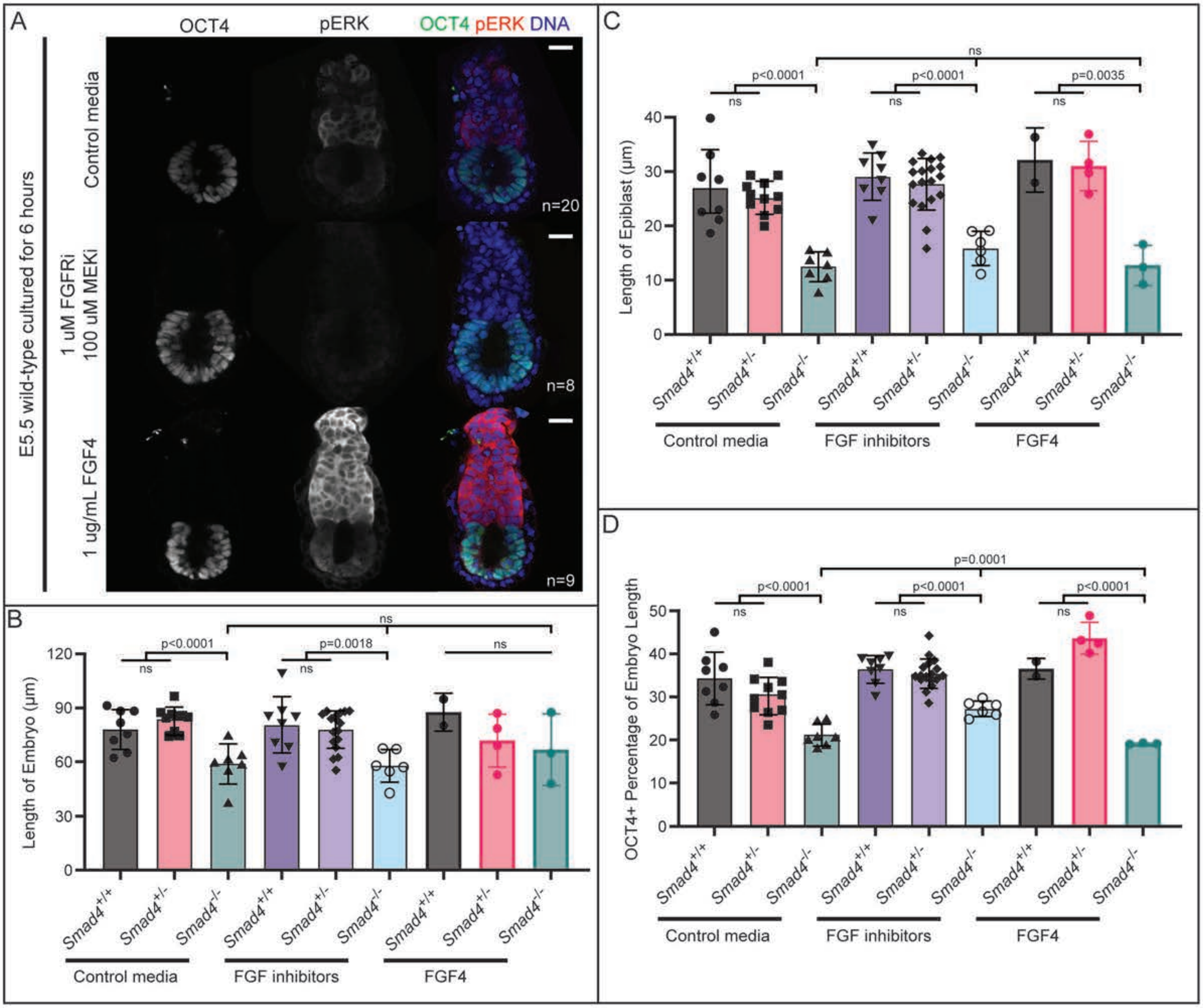
